# Alpha Traveling Waves during Working Memory: Disentangling Bottom-up Gating and Top-down Gain Control

**DOI:** 10.1101/2024.03.21.586061

**Authors:** Yifan Zeng, Paul Sauseng, Andrea Alamia

## Abstract

While previous works established the inhibitory role of alpha oscillations during working memory maintenance, it remains an open question whether such an inhibitory control is a top-down process. Here, we attempted to disentangle this issue by considering the spatio-temporal component of waves in the alpha band, i.e., alpha traveling waves. We reanalyzed two pre-existing and open-access EEG datasets where participants performed lateralized delayed match-to-sample working memory tasks. In the first dataset, the distractor load was manipulated (2, 4, or 6), whereas in the second dataset, the memory span varied between 1, 3, and 6 items. In both datasets, we focused on the propagation of alpha waves on the anterior-posterior axis during the retention period. Our results reveal an increase in alpha-band forward waves as the distractor load increased, but also an increase in forward waves and a decrease in backward waves as the memory set size increased. Notably, our results also showed a lateralization effect: alpha forward waves exhibited a more pronounced increase in the hemisphere contralateral to the distractors, whereas the reduction in backward waves was stronger in the hemisphere contralateral to the targets. In short, the forward waves were regulated by distractors, whereas targets inversely modulated backward waves. Such a dissociation of goal-related and goal-irrelevant physiological signals suggests the co-existence of bottom-up and top-down inhibitory processes: alpha forward waves might convey a gating effect driven by distractor load, while backward waves may represent direct top-down gain control of downstream visual areas.

**Significance Statement:** When exploring the functional role of alpha band neural oscillations during working memory, significant attention has been directed towards its amplitude modulation, with relatively limited exploration of spatial-temporal dynamics of this rather global brain oscillatory signature. The present study seeks to address this gap by examining the directionality of alpha wave propagation during working memory retention. Our findings offer novel insights into the well-established inhibitory role of alpha waves, demonstrating that this function is manifested differently according to their propagation directions: forward waves seem to facilitate bottom-up gating, while backward waves might mediate top-down gain control.

## Introduction

Among all the physiological activation patterns observed with Electroencephalography (EEG), waves in the alpha band (8-12 Hz) are among the most thoroughly studied brain signals. Previous works related alpha band oscillations to inhibitory functions, such as selective spatial attention (Morrow et al., 2023; Peylo et al., 2021; Schneider et al., 2022) or to visual perception (Busch et al., 2009; VanRullen & Macdonald, 2012). Yet, several studies have also demonstrated a connection between alpha waves and working memory. For example, alpha oscillations were persistently observed during working memory retention (see Roux & Uhlhaas, 2014 for review), and causally linked to this process (Chen et al., 2023; Riddle et al., 2020; Sauseng et al., 2009). Moreover, experimental findings revealed that alpha power increased with working memory load (Jensen et al., 2002) and with anticipated distractors (Bonnefond & Jensen, 2012; Payne et al., 2013), suggesting that alpha oscillations may be involved in protecting memory maintenance through an inhibitory mechanism. Other studies have shown that alpha power associated with the gating and inhibition of the processing of irrelevant information is modulated by the perceptual load of the processed stimulus (Gutteling et al., 2022; Jensen, 2023; Noonan et al., 2018). Altogether, these results hint at the crucial role of alpha oscillations in shielding working memory from competing yet task-irrelevant stimuli. However, it remains unclear what are the underlying mechanisms characterizing this process, and whether it involves top-down rather than bottom-up control (Jensen, 2023). One way to address these questions is to consider not only the temporal but also the spatial component of alpha oscillations, that is considering them as traveling waves. These temporospatial dynamics characterize how oscillations propagate through cortical regions, potentially leading to the modulation of excitability across brain areas in a coordinated way.

A growing body of studies has revealed the potential of disentangling distinct functional processes via investigating alpha oscillations as traveling waves propagating in different directions (Aggarwal et al., 2022; Alamia et al., 2020, 2023; Alamia & VanRullen, 2019; Pang et al., 2020). Importantly, the direction of propagation may be associated with different components in distinct cognitive functions. For example, a recent study showed that alpha backward waves (propagating from frontal to posterior regions) were lateralized to the hemisphere contralateral to the unattended visual field, potentially serving as a top-down inhibitory mechanism. By contrast, forward waves (from posterior to occipital regions) increased in the hemisphere contralateral to the attended location, suggesting their relation to visual processing (Alamia et al., 2023). Overall, these studies pave the way for a potential dissociation between distinct candidate mechanisms underlying alpha inhibition in different cognitive functions (i.e., working memory).

Whereas alpha-band traveling waves have been investigated in visual perception and attention, fewer studies explored their relation to working memory processes, although this approach could shed new light on their functional role in working memory. Here we show that considering both the spatial and the temporal component of oscillations can indeed offer novel insight into the inhibitory role of alpha waves in visual working memory. Specifically, we reanalyzed two publicly available EEG datasets that tested visual working memory paradigms. The first dataset manipulated the distractor load during a visual memory task (Feldmann-Wüstefeld & Vogel, 2019), whereas the second dataset varied the set size of memory items (Adam et al., 2018). We took advantage of a method proposed in previous studies (e.g., Alamia & VanRullen, 2019) and computed the normalized power of alpha waves propagating in forward and backward directions. Importantly, our results revealed a dissociation between the distractor load-induced effects, which were predominantly observed in forward waves, and the memory load-induced effects, which were more pronounced in backward waves. We interpret these results as indicating the co-existence of two distinct alpha band phenomena related to inhibitory processes, with forward waves implicated in gating the visual stream, while backward waves appear to contribute to general top-down gain control.

## Methods

### Dataset 1

#### Data Download

Publicly available EEG data reported by Feldmann-Wüstefeld & Vogel (2019) were used. Data were downloaded from https://osf.io/a65xz.

#### Participants

The study by Feldmann-Wüstefeld & Vogel (2019) included three separate experiments. Data from the three experiments included 31 (13 males, M = 21.2 years, *SD* = 2.6; the demographic information of five participants is missing in the dataset), 37 (16 males, *M* = 25.0 years, *SD* = 3.7; the demographic information of five participants is missing in the dataset too), and 36 participants (18 male, M = 22.9 years, *SD* = 3.2) respectively, resulting in a total of 104 volunteers. As pointed out by Feldmann-Wüstefeld & Vogel (2019) data were recorded with the written understanding and consent of each participant.

#### Experimental design and procedure

In each experiment a similar change detection task was used. The task included a memory display consisting of three types of stimuli: colored squares (targets), colored circles (salient distractors), and grey circles (placeholders). Clusters of targets, salient distractors, and two groups of placeholders were presented in one of four different positions (Fig. 1A). In half of the trials, the target cluster was presented in the left or right position, and the salient distractor cluster was presented at the top or bottom (i.e., “targets lateral” condition). In the other half of the trials, the positions of targets and distractors were reversed (i.e., “distractors lateral” condition). While the number of targets was kept constant, a different amount of colored distractors was presented in separate trials: *distractor load* 2 and 4 in experiment 1; *distractor load* 2, 4, and 6 for experiment 2; heterogeneous (4 distractors in 4 different colors) and homogeneous distractors (4 distractors in 2 colors, unused in our study and thus unplotted in Fig. 1A) for experiment 3.

**Figure 1.**
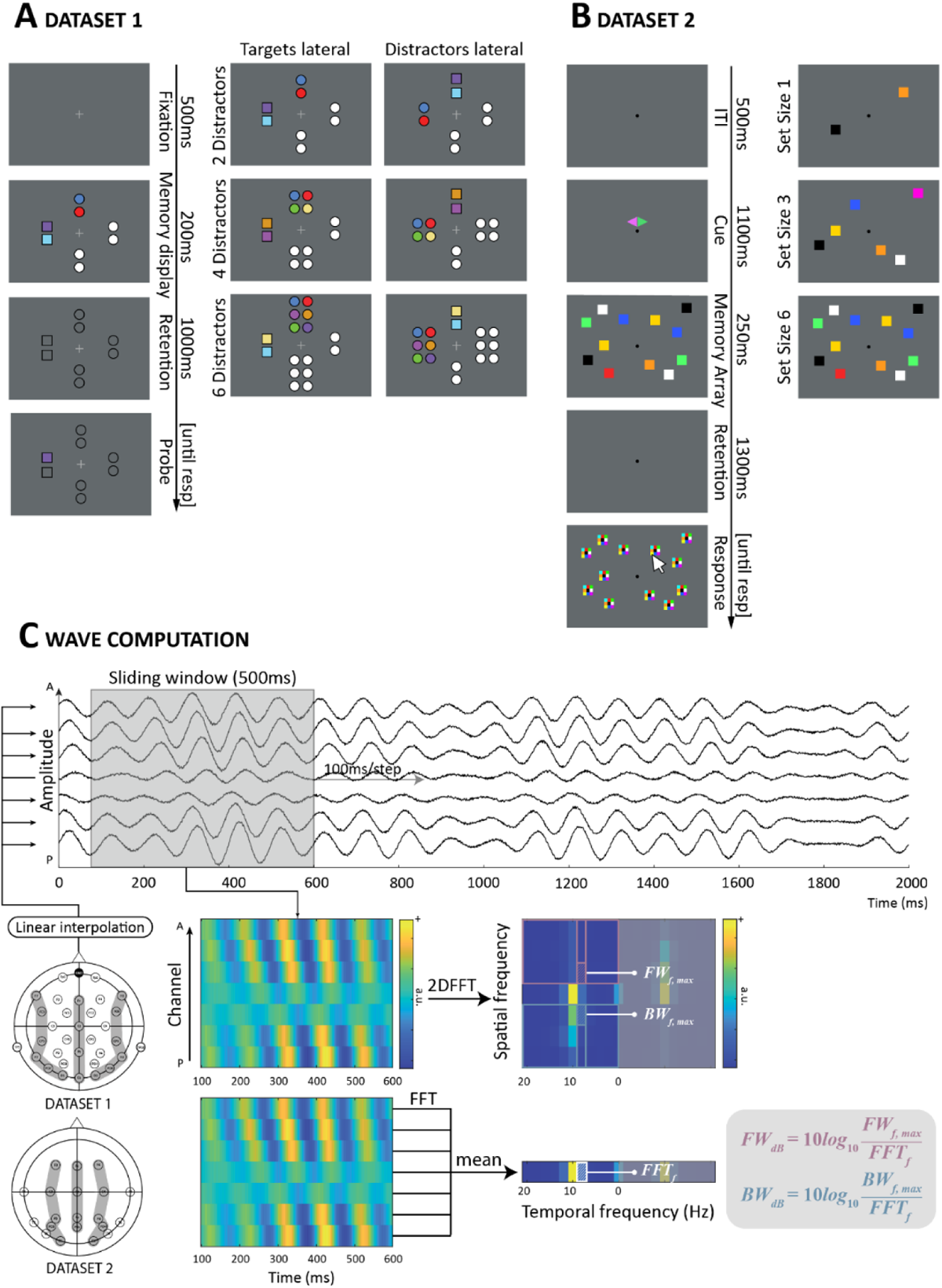
the experimental paradigms and wave computation method. (A) The change detection task of dataset 1 (Feldmann-Wüstefeld & Vogel, 2019). Participants were required to keep in mind the colors of targets (squares) while ignoring the other distracting stimuli (circles). After a short retention interval, participants indicated whether the probe’s color matched with the target presented at the same location. (B) The lateralized whole-report task of dataset 2 (Adam et al., 2018). Participants were instructed to pay attention to and remember the stimuli in the hemifield cued by the green triangle while ignoring the rest. Then, they reported all the colors of the cued stimuli by clicking the color within the matrix corresponding to the items previously presented. (C) Wave quantification method. We selected three electrode axes; midline, left and right hemisphere. The channels within each axis were linearly interpolated into seven channels and sorted according to their spatial location (from posterior to anterior). We employed a 500-ms sliding window at the step of 100 ms, and the resulting segments were fed into two-dimensional fast Fourier Transform. In the power spectrum, the power of forward waves falls into the top-left (or down-right) quarter, whereas that of the backward waves falls into the down-left (or top-right) quarter. For a given temporal frequency and traveling direction, we defined the power as the maximum value in the corresponding column of the respective quarter, normalized by the averaged fast Fourier Transform power of the same temporal frequency. A: anterior, P: posterior, 2DFFT: two-dimensional fast Fourier Transform, FFT: fast Fourier Transform, FW: forward waves, BW: backward waves.

Each trial started with a central fixation cross of 500 ms. Then, the memory display was presented for 200 ms, followed by a retention interval of 1000 ms. A probe followed and was displayed until participants responded. The participant’s task was remembering each target’s color and location while ignoring the distractors. After the probe was presented, they had to indicate whether the color and location of the probe matched the previously presented targets. The trial ended with the participants responding (button press) and a 1000 ms blank screen as the intertrial interval. See Feldmann-Wustefeld & Vogel (2019) for more detail.

#### EEG acquisition and preprocessing

EEG was recorded using a 32-channel BrainProducts actiCap (electrodes positioned according to the International 10/10 System: FP1/2, F7/8, F3/4, Fz, FC5/6, FC1/2, C3/4, Cz, TP9/10, CP5/6, CP1/2, P7/8, P3/4, PO7/8, PO3/4, Pz, O1/2, Oz) with a sampling rate of 1000 Hz. Horizontal and vertical EOGs were recorded. All electrodes were referenced to TP10 and re-referenced offline to the average of all electrodes. For each trial, the dataset provided segmented data from 800ms before to 1600ms after the onset of the memory display. We did not conduct any further preprocessing going beyond the preprocessing already applied in the provided data set.

### Dataset 2

#### Data Download

Publicly available EEG data reported by Adam et al. (2018) were used. Data were downloaded from https://osf.io/8xuk3/.

#### Participants

The second dataset by Adam et al. (2018) consisted of two experiments with 31 (12 female, M = 22.0 years, SD = 3.7) and 45 participants (24 female, M = 21.7 years, SD = 4.2), respectively.

#### Experimental design and procedure

The two experiments from dataset 2 were lateralized whole-report tasks in which participants reported the color of a subset of previously presented targets. The memory display contained two lateralized stimulus clusters, with equal squares on each half of the screen. For each cluster, the squares were randomly distributed within one hemifield, and each square color was chosen without replacement from a pool of nine colors (red, green, blue, yellow, magenta, cyan, orange, white, and black). Repetition of the colors was allowed between hemifields. Before the memory display, a diamond-shaped cue was presented above the central fixation point. The cue was composed of a green and a pink triangle, and participants were instructed to pay attention to the visual hemifield cued by the green triangle. In experiment 1, the number of items in each cluster varied between 1, 3, and 6. In experiment 2, it was kept at a constant level of 6 (Fig. 1B).

Each trial began with a 500-ms fixation point. A small diamond-shaped cue of 1100 ms was then presented above the fixation point. A brief memory display lasting 250 ms was presented, and participants were instructed to maintain fixation on the fixation point while encoding information presented in the visual hemifield indicated by the green side of the cue. The memory display was followed by a retention period of 1300 ms. Participants were required to memorize the colors of each item on the cued side during the retention period, and they reported them by clicking the corresponding color within a 3 × 3 matrix (containing all nine possible colors) presented afterward at each target location. In each trial, the test period lasted until participants had responded to all target items. At last, participants initiated a new trial with an additional mouse click.

#### EEG acquisition and preprocessing

EEG during the whole-report task was recorded using a 20-channel electrodes cap (ElectroCap International, Eaton, OH) from standard 10/20 sites: F3, Fz, F4, T3, C3, Cz, C4, T4, P3, Pz, P4, T5, O1, and O2. Additionally, five nonstandard sites were included: OL (midway between T5 and O1), OR (midway between T6 and O2), PO3 (midway between P3 and OL), PO4 (midway between P4 and OR), and POz (midway between PO3 and PO4). Horizontal and vertical EOGs were recorded to measure horizontal eye movements as well as blinks. All sites were recorded with a right mastoid reference and re-referenced offline to the algebraic average of the left and right mastoids. The sampling frequency was 250 Hz. The dataset contained raw segmented data (from 1400 ms before the onset of the memory display to 1548 ms after the memory display onset). The traveling wave analysis was performed without further preprocessing.

For both datasets, please refer to the original publications by Adam et al. (2018) and Feldmann-Wüstefeld & Vogel (2019) for additional details regarding the presentation apparatus, presentation parameters, pre-test (a change detection task to measure working memory capacity) as well as further information about participants and EEG recording setup.

### Traveling 3Wave Computation

Similar analyses as described by Alamia et al. (2020) were used. Data was analyzed using custom Matlab R2022b (The MathWorks, Natick, MA, USA) scripts and the bayesFactor extension package (Krekelberg, 2023). The data was further exported to JASP for Bayes Factor statistical analysis.

Figure 1C depicts the quantification method of traveling waves. For each dataset, we selected three electrode axes: one in the midline (dataset 1: Fz, Cz, Pz, and Oz; dataset 2: Fz, Cz, Fz, Poz), one over the left hemisphere (dataset 1: F7, FC5, CP5, P7, PO7, O1; dataset 2: F3, C3, P3, PO3, O1), and one over the right hemisphere (dataset 1: F8, FC6, CP6, P8, PO8, O2; dataset 2: F4, C4, P4, PO4, O2). We linearly interpolated the channel array into seven channels to achieve a consistent number of channels across different axes and facilitate the subsequent wave computation. For each trial, we stacked the data of interpolated channels, arranging them from the posterior to anterior regions. Next, a 500-ms sliding window was moved across the data in steps of 100 ms to capture a two-dimensional segment (7 interpolated channels by 500 ms). In the resulting data segments, traveling waves would manifest as planar waves tilted at different angles (i.e., with a monotonic phase shift across channels).

Traveling waves were quantified by the two-dimensional Fast Fourier Transform (2DFFT), which was applied to the data segments considered as images. In the resulting power spectrum of the 2DFFT, the x-axis denotes the zero-centered temporal frequencies, while the y-axis represents the spatial frequencies. Keeping in mind that one of the 2DFFT’s properties is to have conjugation symmetry, we considered the power of forward waves reflected into the top-left (or down-right) quarter, whereas that of the backward waves falls into the down-left (or top-right) quarter. The horizontal midline quantifies the power of standing waves. For a given temporal frequency and traveling direction, we defined the power as the maximum value in the corresponding column of the respective quarter. Yet, as with any other attempts to capture the phasic relationship of oscillations, the traveling wave measure at this stage is susceptible to the temporal oscillatory power, potentially confounding our results. We thus applied a normalization factor using the 1D-FFT power averaged across the seven channels composing each data segment:

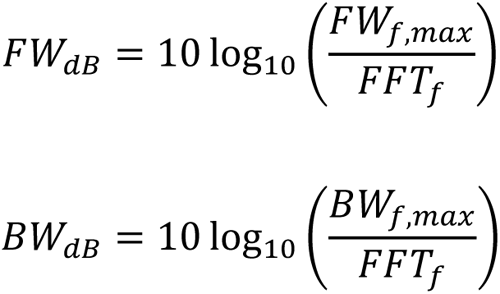

The power in decibels enables a straightforward interpretation of the quantity of traveling waves by comparing them to zero, which serves as a baseline to account for power fluctuations. Subsequently, we computed the power of alpha-band traveling waves by averaging the power of frequencies ranging from 8 to 12 Hz. Furthermore, it is crucial to emphasize that our wave analysis concentrates on the sensor level. This choice is motivated by significant limitations associated with source projections, including the disruption of long-range connections and signal smearing caused by scalp interference (Alexander et al., 2019; Freeman & Barrie, 2000; Nunez, 1974).

### Data analysis

To make the best use of the datasets, we pooled together the sub-experiments within each dataset by their conditions. This leads to a 2 (*lateralized stimuli*: “targets lateral”, “distractors lateral”) by 3 (*distractor load*: “load 2”, “load 4”, and “load 6”) design for dataset 1, and 3 levels of *set size* (“set size 1”, “set size 3” and “set size 6”) for dataset 2.

For each dataset, we conducted three analyses: (1) a Bayesian repeated measures ANOVA with *distractor load* (or *set size* for dataset 2) as the independent variable for midline traveling waves; (2) Bayesian t-tests comparing the power of alpha traveling waves between two lateral axes for each time point; (3) a Bayesian repeated measures ANOVA with the factors of *axes* (i.e., wave power on the axis ipsilateral or contralateral to the items of interest) and *distractor load* (or *set size* for dataset 2) as the independent variables.

Please note the inconsistency of study designs across sub-experiments. For this reason, analyses (1) and (3) did not utilize all the data (see Table 1). For dataset 1, only experiment 2 was included, as it is the sole experiment with three levels of *distractor load*. For dataset 2, only experiment 1 was included, given it is the only experiment with three levels of *set size*.

**Table 1.**
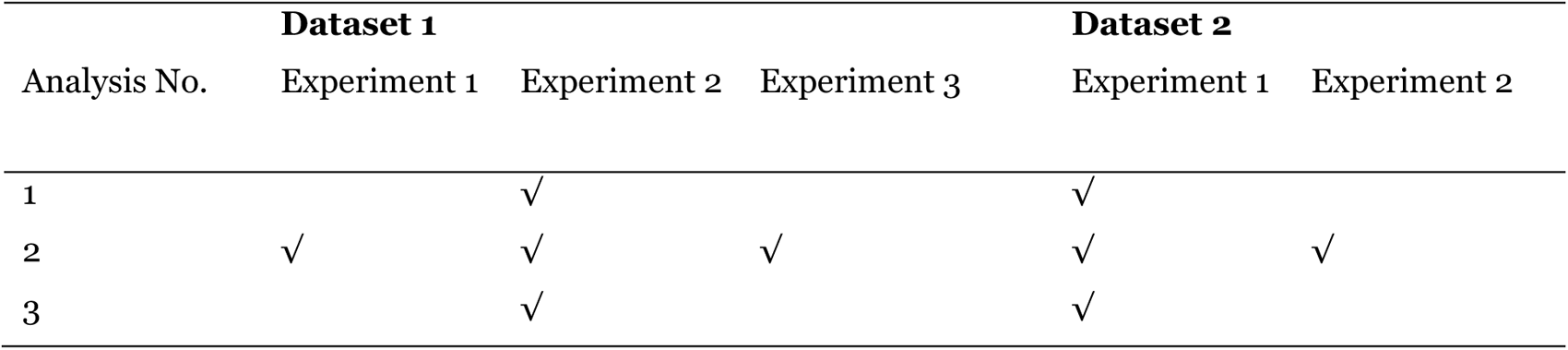
The analyses and utilization of datasets.

## Result

### Midline alpha traveling waves

As the first step, we examined the midline axes to investigate how manipulation of *distractor load* and *set size* modulate alpha traveling waves during the retention period irrespective of lateralization.

In dataset 1, we found that the amount of midline alpha backward waves, but not forward waves, increased when the number of distractors increased (Fig. 2, upper panel). Bayesian repeated measures ANOVA revealed that the observed alpha power of backward waves was more likely under the model with *distractor loa*d than the null model (*BF_incl_* = 24.466). These results suggested that midline alpha backward waves increased when more distractors were presented. Post hoc comparisons indicated substantial evidence for differences between “load 2” and “load 4” (*△M* = -0.024, *_BF10_* = 4.670, *error* = 4.477×10^-7^%) and “load 2” and “load 6” *(△M* = -0.035, *BF_10_* = 13.415, *error* = 7.800×10^-8^%). Between “load 4” and “load 6”, we found indecisive evidence in favor of the null hypothesis (*△M* = -0.011, *BF_10_* = 0.382, *error* = 0.036%). In contrast, we found strong evidence against the effect of *distractor load* on midline alpha forward waves (*BF_incl_* = 0.096).

**Figure 2.**
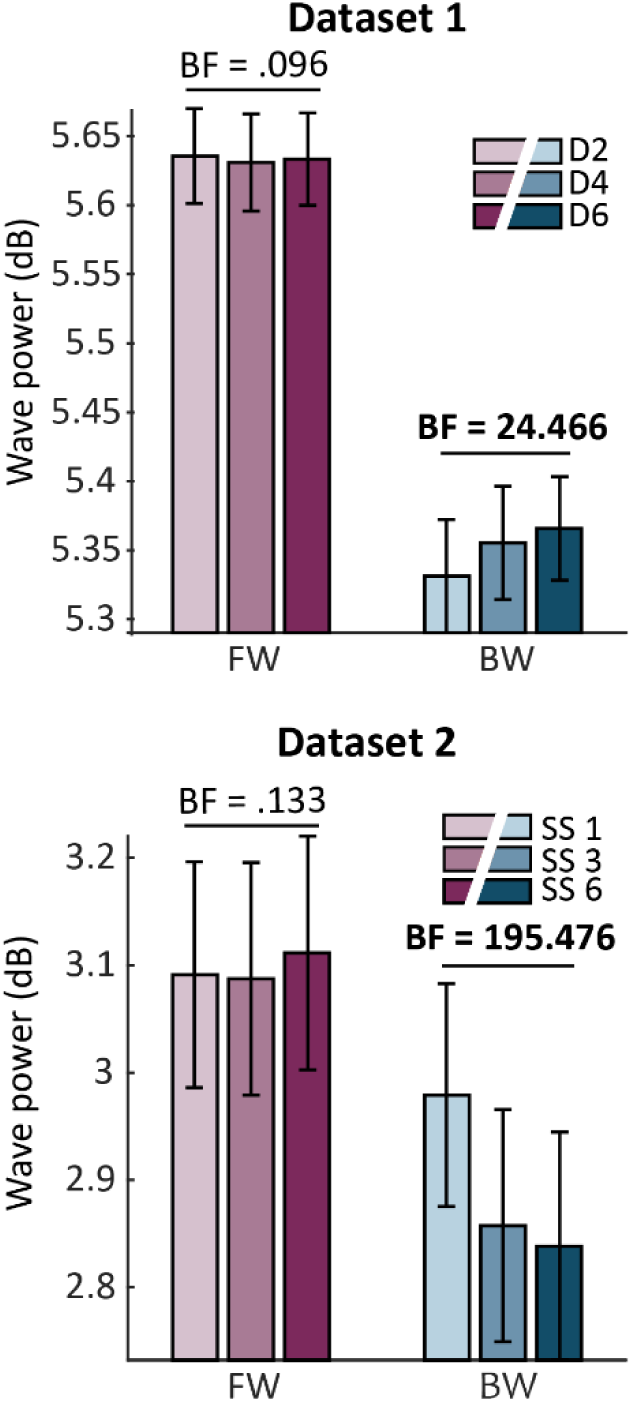
Midline alpha traveling waves during retention periods. Backward waves increased with distractor load in dataset 1 and decreased with set size in dataset 2. BF: the inclusion Bayes factor, D2/D4/D6: distractor load 2, 4, or 6; SS 1/3/6, set size 1, 3, or 6; FW, forward waves; BW, backward waves.

In dataset 2, we also found a modulation of backward waves but not of forward waves; differently than in dataset 1, the midline alpha backward waves decreased with the number of items to be remembered (Fig. 2, lower panel). Our analysis revealed stronger evidence for the effect of *set size* on midline alpha backward waves than the null hypothesis (*BF_incl_* = 195.476); fewer backward waves were found when the *set size* increased. Post hoc comparisons showed significant differences between “set size 1” and “set size 3” (*△M* = 0.122, *BF_10_* = 12.844, *error* = 4.627×10^-8^%), “set size 1” and “set size 6” (*△M* = 0.141, *BF_10_* = 34.020, *error* = 1.673×10^-8^%), but weak evidence against the difference between “set size 3” and “set size 6” (*△M* = 0.019, *BF_10_* = 0.284, *error* = 0.034%). These results indicate that the variation of *set size* also influenced midline alpha backward waves, yet in a different way: in the first dataset, backward waves increased with the number of distractors, whereas in the second dataset, backward waves decreased with the *set size*. As for the midline alpha forward waves, our analysis revealed moderate evidence against the main effect of *set size* (*BF_incl_* = 0.133).

### Lateral alpha traveling waves

Next, we focused our analysis on alpha-band traveling waves propagating in each hemisphere by comparing the power between ipsi- and contra-lateral axes with respect to the stimuli in dataset 1.

We first performed Bayesian t-tests on contralateral vs. ipsilateral wave power in each condition. The line graphs in Fig. 3 depict wave power at each time point. For forward alpha waves, we found a higher power in the axis contralateral to the less attended positions at the later stage of the retention period (1050-1250ms) in both the “target lateral” and “distractor lateral” conditions. By contrast, alpha backward waves were lateralized to the axis ipsilateral to the targets in the “targets lateral” condition, but we found no differences between the two axes in the “distractors lateral” condition.

**Figure 3.**
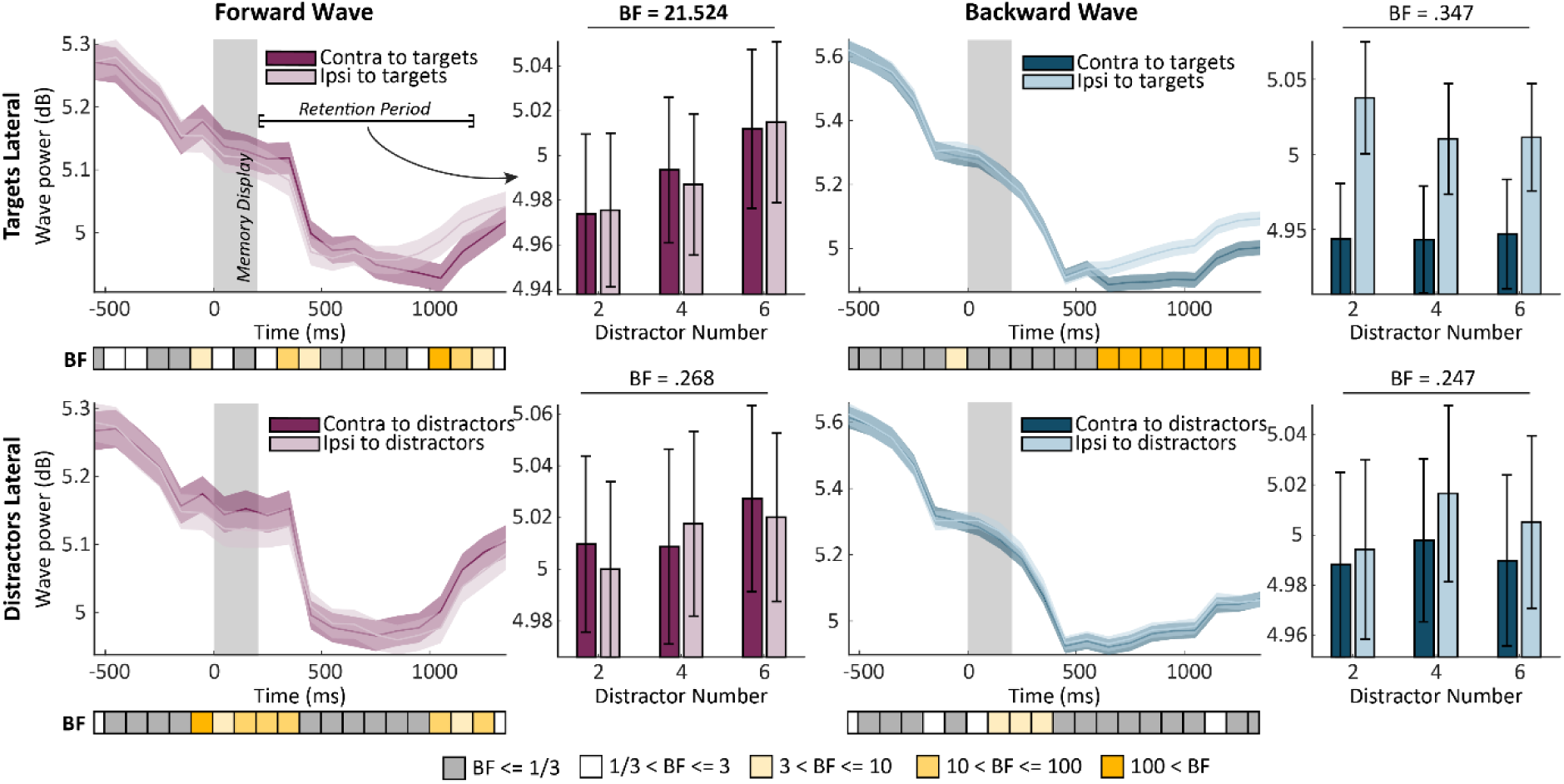
Bilateral alpha traveling waves in dataset 1. Line plots: the power of bilateral alpha traveling waves across time. Below, the color bar indicates the Bayes factors of the power difference between two lateral axes for each time point. Bar plots: the wave power averaged over the retention period. The inclusion Bayes factors above the plots show the main effect of distractor load.

In dataset 2 (the line graphs in Fig. 4), the comparison between the contra-and ipsi-lateral axes revealed similar lateralization patterns regardless of the traveling direction. Specifically, we observed an increase in forward and backward waves in the axis ipsilateral to the target side. Regarding alpha forward waves, this pattern persisted throughout almost the entire analysis window (*BF_10_* > 3), while for backward waves, the effect mainly took place during the cueing (-850 ∼ 50 ms) and the retention period (550 ∼ 1250 ms).

**Figure 4.**
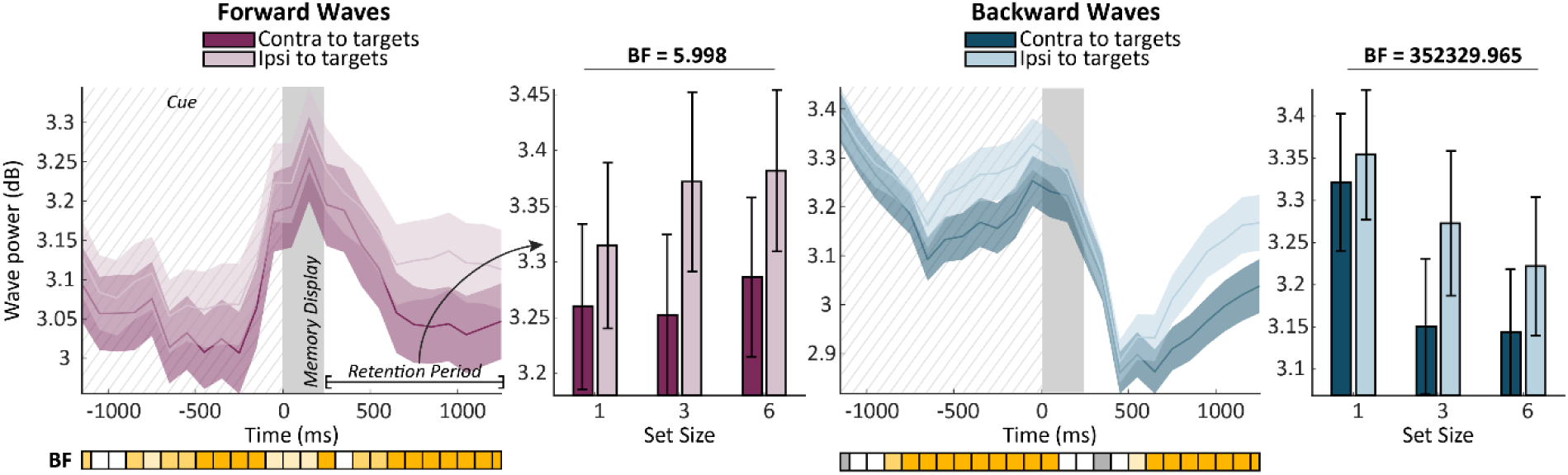
Bilateral alpha traveling waves in dataset 2. The line plots and bar plots represent the bilateral wave power across time and averaged over the retention period, respectively. For more details please refer to the legend of Figure 3.

#### Interaction of axes and effects of distractor load/set size

We then investigated whether the lateralization of traveling waves was modulated by *distractor load* (dataset 1) and the *set size* (dataset 2). The analysis focused on the retention phase by averaging the wave power within these time windows (dataset 1, 200 – 1200 ms; dataset 2, 250 – 1550 ms).

In dataset 1, we conducted four repeated measures ANOVAs for each wave direction and *lateralized stimuli* (the bar plots in Fig. 3). We considered *axes*, *distractor load*, and their interaction term *axes* * *distractor load* as predictors. Overall, the results revealed an increase in alpha forward waves in response to increased *distractor load*, yet only for “targets lateral” condition. Specifically, we found strong evidence for the main effect of *distractor load* (*BF_incl_* = 21.524), whereas the main effect of *axes* and their interaction term remain in favor of the null hypothesis (all *BF_incl_* < 0.173). As for the “distractors lateral” condition, we found no evidence in favor of the alternative models (all *BF_incl_* < 0.268).

We did not observe a similar effect for alpha backward waves. In the “targets lateral” condition, the model considered only *axes* provided the best fit (the analysis of effect showed *axes’ BF_incl_* = 2285.956, whereas for *distractor load BF_incl_* = 0.347 and for *distractor load* * *axes BF_incl_* = 0.835). Regarding the “distractors lateral” condition, none of the effects attained conclusive evidence over the null hypothesis (*axes, BF_incl_* = 0.726; *distractor load, BF_incl_* = 0.247; *axes* * *distractor load, BF_incl_* = 0.069). Overall, these results show that while the amount of alpha forward waves was influenced by *distractor load* in the “targets lateral” condition, this effect didn’t modulate backward waves.

In dataset 2, we conducted two separate repeated measure ANOVAs for each traveling direction, considering *axes*, *set size*, and their interaction *axes * set size* as predictors. The results are shown in the bar plots of Fig. 4. In addition to *axes* (*BF_incl_* = 53885.652), which was already addressed via Bayes t-tests in the section above, we found an increase of alpha forward waves with the *set size* (*BF_incl_* = 5.998). Our analysis also revealed an interaction between *axes* and *set size* (*BF_incl_* = 16.855), suggesting that the increase of alpha forward waves was more prominent on the axes ipsilateral to the attended side.

By contrast, our analysis showed that bilateral backward waves decreased while the *set size* increased. Analysis of effects showed decisive evidence for the main effect of *set size* (*BF_incl_* = 352329.965) and *axes* (*BF_incl_* = 2052.282). Moreover, the interaction between the two factors provided strong evidence (*BF_incl_* = 24.979), indicating that the decrease of backward waves was more substantial in the axis contralateral to the target position. All in all, these results showed that alpha forward waves increased when the overall number of items on the unattended side increased. In contrast, backward waves decreased with the number of items on the attended side.

## Discussion

The present study aims to investigate the functional roles played by forward and backward alpha-band traveling waves during working memory tasks. Specifically, we showed how different features manipulating working memory processes, namely distractor loads (i.e., number of distractors, dataset 1; Feldmann-Wüstefeld & Vogel, 2019) and set size (i.e., number of targets, dataset 2; Adam et al., 2018), differentially modulate forward and backward waves. Fig. 5 summarizes these results schematically. In dataset 1, the alpha power of midline backward waves (unplotted) and the power of bilateral forward waves increased with the distractors’ load during the retention phase. In addition, we showed that alpha backward waves, but not forward waves, were lateralized to the axis ipsilateral to the encoded targets. In dataset 2, we found that larger set size led to an increase in the propagation of bilateral forward waves and, conversely, a decrease of backward waves overall. These effects interacted with the hemispheres where the waves propagate: the increase of forward waves was more prominent on the axis contralateral to the distractors, whereas the reduction of backward waves was more pronounced on the axis contralateral to the targets. The overall traveling wave amount, however, was higher on the axis contralateral to the distractors for both wave direction. In the following, we interpret our results in light of the differential roles of forward and backward alpha-band waves during visual working memory tasks.

**Figure 5.**
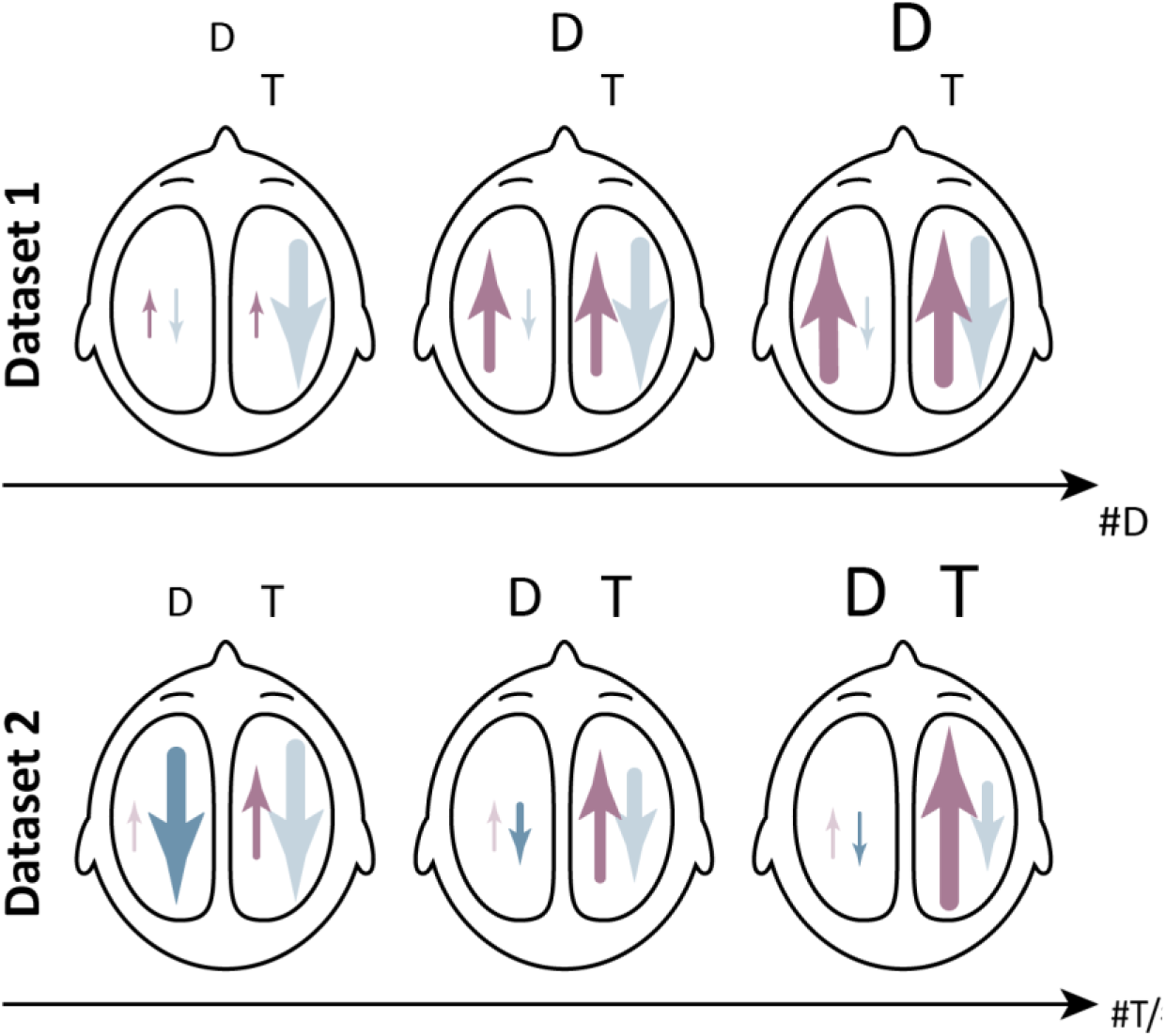
Schematic summary of the results. The size of the arrows indicates the power of traveling waves. We denote the effects of interests with less transparency. D: distractors, #D: distractor load, T: targets, #T; set size.

### Alpha forward waves reflect gating during retention

Our findings in alpha forward waves go in line with the idea of alpha waves inhibiting irrelevant information by gating mechanism (Jensen & Mazaheri, 2010). Specifically, in dataset 1, we observed an increase in forward waves accompanying an increase in the central distractor load; assuming that stronger forward alpha waves represent gating of information flow from lower to higher visual areas, a higher distractor load would also require more of such gating. Accordingly, in dataset 2, there was a similar effect observed for forward waves contralateral to distractors: an increase in the number of distractors went hand in hand with increasing forward waves, indicating more rigorous gating in the hemisphere contralateral to distractors. No such effect was observed, however, for the hemisphere contralateral to targets. As no targets should be filtered away, even at high target load, there was no increase of forward waves obtained in the contralateral hemisphere.

Such gating of sensory information (Jensen & Mazaheri, 2010) may not be under direct top-down control (Jensen, 2023), also indicated by the ‘forward’ traveling direction in the present study. It has been suggested that the rhythmic fluctuation of alpha oscillations provides pulsed inhibitions to gate the flow of sensory input (Jensen & Mazaheri, 2010). As a result, those that fall into the excitatory phases (i.e., the duty cycle) get better processed than those that run into the phases of inhibition. Most likely, the gating of information does not reside in early visual areas but instead somewhere slightly higher in the visual hierarchy (Jensen, 2023; Peylo et al., 2021). In contrast to that, here, we suggest that gating does not take place at only one level within the visual hierarchy; instead, we proposed that this gating mechanism be implemented by alpha forward waves, superimposing the low-level visual input and carrying out real-time screening along the information stream while it flows toward higher regions.

#### Distinct roles for stimuli-evoked and inhibitory alpha-band forward waves

However, it is worth noting that alpha-band forward waves may play distinct functional roles during different stages of the current working memory task (e.g., during memory display vs. retention phase). A recent study suggested that alpha-band forward waves may appear during low-level visual stimulation, as they reported forward waves during visual stimulation but less so in the absence of sensory input (Pang et al., 2020). Similarly, during presentation of the memory display (see Fig. 3) we observed more contralateral than ipsilateral forward waves in both “target lateral” and “distractor lateral” conditions, in line with the hypothesis that forward waves reflect sensory stimulations; as both targets and salient distractors were perceptually more salient than the placeholders. In other words, the colorful stimuli introduced more low-level visual information compared to the grey placeholder on the other side of the screen, which may lead to more alpha forward waves. However, more visual perception-related forward alpha waves might be difficult to dissociate from more memory-related forward alpha waves as we see them during the retention interval. In the current task they are observed during different task phases, though, and they respond differently to changes in target and distractor load. Possibly, more fine grained spatial analysis (like suggested by Zhang et al., 2018) of traveling waves could prove successful for dissociating those functionally different waves in the future.

### Alpha backward waves reflect top-down gain control

In contrast to forward waves, alpha backwards waves were not modulated by distractor load. Instead, backward waves were generally strong in the hemisphere ipsilateral to targets. This might indicate an inhibitory function of alpha backwards waves in working memory, similar to effects obtained on spatial attention (Alamia et al., 2023). Moreover, this pattern is reminiscent of findings reported on stationary posterior alpha amplitude when spatial attention is shifted to one visual hemifield, resulting in increased alpha activity ipsilateral to the locus of attention (Foxe & Snyder, 2011; Medendorp et al., 2007; Sauseng et al., 2005; Thut et al., 2006; Worden et al., 2000).

In the hemisphere contralateral to targets, backward alpha waves are inversely related to target load. An increased number of targets led to a reduction of backward waves (Fig. 5). Considering alpha backward waves as an inhibitory top-down biasing signal here, a reduction of backward waves with increasing number of targets to retain would lead to release of inhibition in more down-stream, lower visual areas. Therefore, backwards alpha waves in the here reported data might be a correlate of inhibitory gain control in the visual system.

### Linking past studies of alpha traveling waves

While a substantial body of literature has firmly established the inhibitory role of alpha waves during working memory performance (Bonnefond & Jensen, 2012; Gutteling et al., 2022; Jensen, 2023; Jensen & Mazaheri, 2010; Khader et al., 2010; Klimesch et al., 2007; Riddle et al., 2020; Sauseng et al., 2009; Wianda & Ross, 2019), the notion of alpha waves as traveling waves in the human neocortex has only recently been fully considered (Soroka & Idiart, 2021; Zhang et al., 2018). In line with our results, these recent studies have suggested alpha waves traveling at different propagation directions during the different stages of a memory task; not only within the prefrontal cortex (Bhattacharya et al., 2022) but also on a large scale (Mohan et al., 2022 providing evidence for the preferred traveling axis to be posterior-to-anterior). To our knowledge, the current study is the first one to investigate alpha traveling waves with respect to the load of relevant/distracting information during the retention period of working memory tasks. Appling the approach reported in the current study to retro-cuing visual working memory tasks could help investigating if alpha traveling waves may be involved in erasing already stored information, as suggested by Soroka & Idiart (2021). Overall, our results supplemented the current understanding of the link between large scale alpha traveling waves and working memory.

## Conclusion

Together, our study may provide neural evidence for the co-existence of two potential inhibitory mechanisms of alpha waves for the first time (Jensen, 2023). These two mechanisms may reside in alpha waves, which propagate in opposite directions along the posterior-anterior axis: alpha backward waves may represent a direct top-down biasing signal for gain control, whereas forward waves might convey a bottom-up gating effect.

One of the main conclusions of this work is the importance of considering not only the temporal but also the spatial dynamics when investigating neural oscillations. For example, considering only the power difference between the target- and distractor-induced effect might not capture the true complexity of how targets and distractors are processed in working memory. Similarly, we presume that considering oscillations as traveling waves propagating in different directions could shed light on some of the contradicting studies investigating the functional roles of alpha waves in working memory.

## Acknowledgment

This project was funded by the European Union under the European Union’s Horizon 2020 research and innovation program (grant agreements No. 101075930 to Andrea Alamia). The copyright holder for this those of the author(s) only and do not necessarily reflect those of the European Union or the European Research Council (ERC). Neither the European Union nor the granting authority can be held responsible for them.

